# 2-bromo-2’5’-dihydroxychalcone analogue Inhibits Endothelial Migration by Targeting VEGF-induced ERK 1/2 Phosphorylation

**DOI:** 10.1101/2023.09.11.557154

**Authors:** Aamir Hussain, Joseph Festa, Harprit Singh

**Author notes:** Corresponding author: Aamir Hussain, Leicester School of Allied Health Sciences, De Montfort University, The Gateway, Leicester LE1 9BH, UK Email.

## Abstract

Angiogenesis, the process of new blood vessel formation, is characterized by three essential hallmarks: endothelial proliferation, migration, and differentiation. Each is integral in angiogenesis related diseases, especially cancer. With drug efficacy stagnated due to acquired drug resistance and off target side effects, the need for combinatorial therapy is ever more present. To identify new compounds that could aid current antiangiogenic therapies, we report the preliminary mechanistic evaluation of a 2-bromo-2’5’-dihydroxychalcone analogue and its antimigratory effects on endothelial cells. After the synthesis and validation of the 2-bromo-2’5’-dihydroxychalcone analogue (AH9), its effect was tested in vitro using human umbilical vein endothelial cells (HUVEC). Initial investigations into 2-bromo-2’5’-dihydroxychalcone effect in vitro was conducted with a cell proliferation assay including MTT, afterward endothelial migration was measured with the scratch assay in subsequent functional studies. For mechanistic evaluation, vascular endothelial growth factor (VEGF) induced ERK phosphorylation using western blot was implemented. AH9 inhibited VEGF-induced ERK ½ phosphorylation similar to that of known antiangiogenic drug Sorafenib at all three concentrations 100 μM (46%, *p* = 0.003), 30 μM (64%, *p* = 0.0002) and 10 μM (91%, *p* = 0.0001). In a scratch assay model, whilst sorafenib at 3 μM was not able to limit migration after 8-hr compared to an untreated control (p = 0.0978), AH9 did (17.41%, p = 0.0079). Furthermore, AH9 was able to inhibit ERK ½ phosphorylation in a concentration dependent manner 100 μM (46%, p = 0.003), 30 μM (64%, p = 0.0002) and 10 μM (91%, p = 0.0001) compared to the VEGF control. These preliminary findings support that AH9 could be exerting antimigratory effects through the inhibition of the VEGF induced MAPK/ERK pathway. This forms the foundation for further studies to explore chalcone analogues in hope to aid current antiangiogenic therapeutic strategies as potential angiogenic inhibitors.

## Introduction

The significance of angiogenesis in tumour growth and development has made it a prime target for anti-cancer research. Over the years, numerous strategies have been hypothesized to disrupt this process with the goal of depriving tumours of their blood supply and halt neoplastic cell proliferation (Ferrara and Kerbel, 2005; Folkman, 2006; Mahar et al., 2009; Gotink and Verheul, 2010; Ebos and Kerbel, 2011; Vasudev and Reynolds, 2014). Angiogenesis itself is a highly regulated multi-faceted process, both in physiological and pathological settings. Three hallmarks encapsulate the process: endothelial cell proliferation, migration, and differentiation. Each of these sub processes utilises a multitude of signalling molecules to coordinate and regulate angiogenesis. Signalling pathways such as angiopoietin-TIE2, MAPK/ERK and VEGF family of ligands and receptors are heavily involved as targets for tumour focused antiangiogenic therapy (Sullivan and Bicknell, 2003). ERK ½ kinases, are key mediators in the signalling cascade to trigger endothelial cell proliferation and migration. The MAPK/ERK pathway contributes to a whole host of cellular processes including regulating T cell activation, phosphorylation of the p53 transcription factor and activation of phospholipase A2 (PLA2) in mast cells (Shaul and Seger, 2007). Regulation of this pathway is key to prevent tumorigenesis, as overexpression of ERK½ can lead to inhibition of apoptosis. In turn this can lead to over proliferation of cells and as a result tumour development.

Current antiangiogenic approaches are centred in either preventing, disrupting, or regressing neo vasculature formation (Samant and Shevde, 2011; Al-Abd et al., 2017). A common approach to block key receptors such as VEGFR-2 has yielded small receptor tyrosine kinase inhibitors (sRTKI’s) like Sorafenib and Sunitinib. However, their theoretical efficacy has not translated to improved clinical outcomes (Bergers and Hanahan, 2008). Within this, evasive and acquire resistance has further stunted the therapeutic use of these antiangiogenic drugs to halt tumour growth in a clinical setting (Bergers and Hanahan, 2008). Hence, there arises a demand not only for novel antiangiogenic drug candidates but also for targeting the secondary processes within the pathological vasculature to holistically inhibit tumour-induced angiogenesis (Loges, Schmidt and Carmeliet, 2010; Ebos and Kerbel, 2011). One strategy is the use of small chemical scaffolds. Chalcones also known as diphenylpropenones are a group of diaryl compounds that situated within the umbrella group of polyphenols found in nature. They bare similar structures to other known phenols such as flavonoids, stilbenes and anthocyanidins (Mirossay et al, 2017). Chemical scaffolds such as chalcones have shown to exhibit antiangiogenic activity via downregulation of VEGFR-2 and signalling mediators such as ERK ½ (Mojzis et al., 2008; Lee et al., 2012; Mahapatra, Bharti and Asati, 2015). Recent studies have shed light on new chalcones derivatives that have been shown to inhibit the expression of ERK ½. Rizeq and colleagues reported novel nitrogen based chalcones capable of inhibiting angiogenesis in chorioallantoic membrane model (CAM) via targeting JNK 1/2/3 and ERK ½ (Rizeq et al, 2021). Kuruc and colleagues identified antimigratory activity of a novel phenolic chalcone L1 against Human cervical carcinoma cells (HeLa) and attributed the modulation of Akt, JNK, p38 and ERK ½ to the activity observed (Kuruc et al, 2021).

Nam et al (2003) discovered the unexpected antiangiogenic activity of 2-chloro-2’5’-dihydroxychalcone. Different analogues of the 2’5’-dihydroxychalcone scaffold were explored but only 2-chloro-2’5’-dihydroxychalcone showed antiangiogenic activity, far superior to the other derivatives. To explore other analogues, we report in this study the functional and mechanistic capabilities of a novel brominated chalcone scaffold, 2-bromo-2’5’-dihydroxychalcone (Figure 1) against HUVEC proliferation and migration, using Sorafenib as a reference control.

## Methods

### Reagents

Antibodies against, phospho-ERK1/2 (CAT: #4377S), ERK1/2 (CAT: #9102S), β-actin (CAT: #8457S) including the IgG secondary antibody (CAT: #7074P2) were purchased from Cell Signalling Tech. VEGF ligand (recombinant human protein) was ordered from ThermoFisher (CAT: #PHC9391).

### Synthesis of 2-bromo-2’5’-dihydroxychalcone (AH9)

2’5’-Dihydroxyacetophenone (6.57mmol) and pyridinium p-toulene sulphonate (0.159mmol) were dissolved in anhydrous dichloromethane and stirred for 30 minutes at 37°C. A solution of 3,4-dihydro-2H-pyran (37mmol) in dichloromethane was added and stirring continued for 4h. The solution was washed with water and the organic layer dried over anhydrous magnesium sulphate. Removal of the solvent under vacuo yielded crude 2’5’-bis (tetrahydropyran-2-yloxy) acetophenone as a straw coloured oil.

Barium hydroxide octahydrate (6.57mmol) was added to a stirred solution of the appropriate benzaldehyde (6.57mmol) and crude 2’5’-bis(tetrahydro-pyran-2-yloxy)acetophenone at room temperature. After 18h the reaction mixture was washed with water and the organic layer dried over anhydrous magnesium sulphate. Removal of the solvent in vacuo yielded crude (E)-1-[2,5-(bis(tetrahydropyran-2-yloxy)-phenyl]-3-phenyl-2-propene-1-one as an orange solid.

A solution of p-toulenesulphonic acid (2.34mmol) and crude (E)-1-[2’5’-(bis(tetrahydropyran-2-yloxy)-phenyl]-3-phenyl-2-propene-1-one in methanol was stirred at room temperature for 30mins. The reaction mixture was quenched with water and extracted with ethyl acetate. The organic extracts were dried over anhydrous magnesium sulphate and the solvent removed under vacuum. The residue was purified by either flash column chromatography (hexane/ethyl acetate 8:2) or recrystalisation from ethanol. (E)-1-(2,5-dihydroxyphenyl)-1-(2-bromophenyl)-2-propene-1-one (AH9).

## Analytical confirmation of AH9 synthesis

Nuclear magnetic resonance spectroscopy (NMR) was used as the primary structure confirmation tool. 1H and 13C NMR spectra were recorded on a Bruker Avance super-conducting AV400 NMR spectrometer at 30°C. Chemical shifts are reported in units relative to the TMS signal standard and coupling constants (J) are expressed as Hertz (Hz). 1H NMR peak information is provided in the following format: number of protons, multiplicity, coupling constant (where appropriate) and peak signal. Multiplicities are reported as: singlet (s), doublet (d), triplet (t), doublet of doublets (dd), doublet of doublet of doublets (ddd), doublet of triplets (dt), triplet of doublets (td), multiplet (m).

High resolution mass spectrometry (HRMS) was carried out using a Thermo Scientific LTQ Orbritrap XL mass spectrometer at EPSRC UK National Mass Spectrometry Facility, Swansea University. Infra-red (IR) spectra were recorded on a Bruker Alpha ATR module (101873) spectrophotometer with bond absorptions reported as cm-1.

Recrystalisation from ethanol. Red/orange crystals (35%). mp 177-179oC; TLC: Rf = 0.13 (hexane/ethyl acetate 8:2); IR: (ATR)/cm-1 3354 (OH), 1641 (C=O); 1H NMR (ppm, DMSO-d6): 6.85 (1H, d, J = 9.4Hz, Ar), 7.04 (1H, dd, J = 9.4, 3.1Hz, Ar), 7.39 (1H, ddd, J = 7.8, 1.9Hz Ar), 7.48 (2H, m, Ar), 7.75 (1H, dd, J = 9.4, 1.4Hz, Ar), 7.91 (1H, J = 15.6Hz, C=CH), 8.02 (1H, J = 15.6Hz, C=CH), 8.14 (1H, dd, J = 9.4, 1.4Hz Ar), 9.20 (1H, s, OH), 11.50 (1H, s, OH); 13C NMR (DMSO-d6) 115.1, 118.3, 121.3, 124.5, 125.4, 125.5, 128.8, 132.3, 133.3, 133.8, 141.1, 149.6, 154.2, 192.6; HMRS found [M+1]+ 318.9969, C15H11O3Br [M+1]+ 318.9964; Purity > 99% HPLC method 1 (Retention time 4.114 min).

### MTT assay

Primary Human Umbilical Vein Endothelial Cells (HUVEC) were acquired from Gibco Invitrogen and cultured in medium 200 supplemented with 10% foetal bovine serum (FBS) and 2% endothelial growth supplement. All experiments were performed between cell passage 3-6. HUVECs were seeded at a density of 3.5 x10^3^ per well in a 96 well plate overnight. Cells were treated with different concentrations of AH9 for 72-hr to form final set of concentrations between 10 μM to 100 nM in media. Cell proliferation was measured via Absorbance of each well was at 560 nM using a Promega GloMax Discovery system, data was exported to Microsoft Excel and analysed to form % control cell viabilities. Each assay was performed 4 times for preliminary screens and 5-6 times for EC/IC50 determinations.

### Scratch (Wound Healing) Assay

HUVECs were seeded at 500,000 cells/well in a 6 well plate and treated with different concentrations of AH9 and Sorafenib in the presence of 2% FBS. Cells were cultured to 90% confluency before being scratched down the middle by a 1 mL pipette tip. Images were taken at time points (0-hr, 4-hr and 8-hr) using the EVOS FL Cell Imaging System (magnification, X100) and the average distance between the wound was calculated over the time points as a measure of migration. Three images were taken at 3 randomly selected fields within each well and were statistically analysed from three independent experiments.

### Cell Stimulation and Western Blotting

HUVECs were seeded at 500,000 cells/well and grown in LSGS media overnight. AH9 was added at 100 μM 30 μM and 10 μM for 1 hr with or without VEGF (30 ng/ml) for 15 minutes. Media was then removed, and each well washed with cold PBS. Total proteins were extracted using laemelli buffer (2X) containing sodium orthovanadate (80mM) and dithiothreitol (216mM). Proteins were separated using gel electrophoresis and transferred onto nitrocellulose membranes using the iBlot™ 2 dry blotting system. The membrane was blocked using 5% non-fat dried milk powder dissolved in TBS-Tx100 (TBS-Tx100: Tris (5mM), NaCl (15mM), 0.05% Tween 20, Triton 100x (20%) pH adjusted to 7.5 using HCL). Membranes were incubated with relevant primary and secondary antibodies (p-ERK ½, ERK ½, β-actin at 1:1000 dilution with Anti-rabbit IgG secondary antibody at 1:2000 dilution). Membranes were developed using ECL and densitometry of protein bands performed using Image J. Selected antibodies were used to probe for desired proteins. Levels of proteins were quantified and normalised using β actin as an internal control. Ratio of phosphorylated (activated) protein vs total protein was expressed as means ± SEM.

### Statistical analysis

Experiments were conducted in triplicates and repeated at least three times. Ordinary one-way ANOVA analysis was conducted using Microsoft Excel STAT package to analyse the MTT assay. For the scratch assay and western blots, a multiple comparative analysis was conducted using the Tukey-Kramer multiple comparisons post hoc unpaired student t-test via GraphPad Prism 7.0b software.

## Results

### AH9 inhibits HUVEC proliferation

Endothelial cell proliferation is one of the key hallmarks of the angiogenesis cascade. To assess the preliminary effects of AH9 on HUVECs, we utilised the MTT assay to quantify the inhibition of endothelial cell proliferation. AH9 was able to inhibit HUVEC proliferation in a dose dependent manner. EC50 value after 72 hr exposure of AH9 and Sorafenib was able to limit 50% of HUVEC proliferation at 4.01 μM (Figure 1D).

### AH9 inhibits HUVEC migration

Building on from endothelial cell proliferation, we looked to observe the anti-migratory activity of AH9. To examine this, we performed the wound healing assay to compare the anti-migratory effects of AH9 vs Sorafenib over an 8-hr time course. Both AH9 and Sorafenib at 10 μM were too concentrated to examine anti-migratory activity hence a lower concentration of 3 μM was used as shown in (Figure 2A).

Interestingly, results for AH9 at 3 μM demonstrated a significant increase in anti-migratory activity at 4-hr when compared to Sorafenib, with AH9 appearing to show regression in HUVEC migration (when both were compared to their time 0hr controls); (at 4-hr, AH9 vs Sorafenib = (−4.9%) vs (34.5%), respectively, p = 0.0115); whereas no significant change was observed at 8-hr (at 8-hr, AH9 vs Sorafenib = (16.2%) vs (52.7%), respectively, p = 0.2602). Figure 2C, demonstrates the width of wounds for each experiment clearing presenting a difference between wounds size for AH9 3 μM (17.41%, p = 0.0079) and Sorafenib 3 μM (49.5%, p = 0.0978) compared to the untreated control over the 8-hr period (80%, p = 0.0027). To confirm, significance in migration was observed for the untreated control at both 4 and 8-hr (55%, p < 0.01, 80% p < 0.01) as seen in (Figure 2B).

### AH9 inhibits VEGF-induced ERK1/2 phosphorylation

AH9 was able to significantly inhibit ERK ½ phosphorylation at all three concentrations 100 μM (46%, p = 0.003), 30 μM (64%, p = 0.0002) and 10 μM (91%, p = 0.0001) compared to the VEGF control. Sorafenib at 30 μM also showed complete inhibition of ERK ½ phosphorylation (100%, p < 0.0001) with no observable signal when compared to the rest of the bands on the nitrocellulose membrane. Unexpectedly less inhibition was seen at the higher concentrations (100 μM and 30 μM) when compared to 10 μM results (Figure 3).

## Discussion

Identifying chemical scaffolds with promising antiangiogenic activity to complement existing antiangiogenic therapies is a needed endeavour. In this study our preliminary data suggests that AH9 exhibits similar anti-proliferative activity at micromolar concentrations against HUVECs as does positive control sorafenib. But more importantly, AH9 exhibited significantly more potent anti-migratory effects against HUVECs than Sorafenib in this *in vitro* model at 8-hr vs matched untreated control. The results demonstrate that AH9 has strong inhibitory effects on endothelial cell proliferation and potent effects on endothelial migration. These are two hallmark characteristics of an agent capable of being developed as an angiogenesis inhibitor, which is why effects on endothelial cell proliferation and migration were explored. Mechanistic evaluation identified that AH9 could be exhibiting its anti-proliferative and anti-migratory activity via the inhibition of VEGF-induced phosphorylation of ERK which was seen at all three concentrations 100 μM (46%, p = 0.003), 30 μM (64%, p = 0.0002) and 10 μM (91%, p = 0.0001). ERK signalling is integral in the angiogenesis cascade (Srinivasan et al, 2009). Many antiangiogenic compounds have shown to downregulate ERK pathway to blunt endothelial cell proliferation and migration. Considering the involvement of ERK kinases in numerous signalling pathways, the inclusion of angiogenic inhibitors that disrupt ERK signalling proves to be a valuable addition to a long-term antiangiogenic regimen. Murphy and colleagues highlighted the significance of MAPK/ERK focused angiogenesis inhibition in sorafenib’s anti-tumour effects (Murphy et al., 2006). Zhang and colleagues put forward the case that p-ERK could be even viewed as a potential biomarker for Sorafenib sensitivity. Owing to the key importance p-ERK plays in the mechanism of action for Sorafenib (Zhang et al, 2009). Recently, Takao and colleagues also observed this where the oncogenic properties of endometrial cancer stem cells were inhibited by Sorafenib via the MAPK/ERK pathway (Takao et al, 2022). As Sorafenib works via this MAPK/ERK pathway, owing much of the anti-tumour effects to blunting ERK 1/2 effects this supports the notion of AH9’s ability to potentially exert antiangiogenic effects similar to that of a known licensed antiangiogenic drug.

These results also link back to original work carried out by Nam et al (2003) on the antiangiogenic activity of 2-chloro-2’5’-dihydroxychalcone. They reported sub micromolar inhibition of HUVEC proliferation (Ic50 0.03 ng/mL) and *in vivo* inhibition of tumour growth by 60.5% in BDF1 mice bearing Lewis lung carcinoma cells at 50mg/kg/day. The data also complies with the work of Varinska et al on Q797 (4-hydroxychalcone) (Varinska et al, 2012).

They concluded that at 100 μM Q797 was able to inhibit VEGF-induced ERK-phosphorylation. Here we report a 10-fold lower and much safer concentration (10 μM) to inhibit ERK phosphorylation (p < 0.0001).

### Limitations

At the higher concentrations (100 μM and 30 μM) inhibition of ERK phosphorylation was significantly less 10 μM, which could be likely due to poor solubility in serum free media. The 5% serum media used for the anti-proliferative work was tolerated up to concentrations of 30 μM. With 10 μM being further diluted before being administered into the well, we believe its solubility was more favoured and as a result it was able to exhibit the inhibitory activity seen in Figure 3A. Intracellular over saturation of AH9 could also be the reason behind the reduced ERK inhibition when compared to a more commonly seen dose (10 μM) in ligand binding studies of lead compounds.

### Future directions

To summarise, these results provide an introductory understanding about the molecular mechanism of action for AH9; in relation to its anti-migratory effect against HUVECs. ERK function is a fundamental signalling molecule for endothelial proliferation and migration. These results show the potential of AH9 as a potent VEGF induced ERK phosphorylation inhibitor. The findings also relate to the significant inhibition of ERK phosphorylation by Sorafenib conducted in this study (p < 0.0001). As highlighted earlier these outcomes support the literature studies on Sorafenib’s effects on ERK expression. But they also further demonstrate how AH9, a simple molecular compound, exhibited similar activities to that of a licensed FDA approved drug for antiangiogenic therapy, on reducing ERK phosphorylation *in vitro*.

Further work is needed to investigate the inhibitory effect of AH9 on the VEGF/VEGFR2/MAPK/ERK signalling pathway. Especially to further investigating specific targets within endothelial migration Such as specific NOTCH transcription targets or PI3K, PDK1 and p38 MAPK all of which are downstream signalling mediators from VEGF-induced VEGFR-2 activation, to pinpoint the molecular mechanism of action by which AH9 inhibited HUVEC migration (Lamalice, Le Boeuf and Huot, 2007). These needs to be followed by VEGFR2 siRNA gene silencing experiments to see if AH9 can lose its ERK inhibitory ability in culture cell model. This should be followed by developing VEGFR-2 knockout mice in a murine model to assess the changes or selectivity of AH9’s effects in an *in vivo* setting.

## Conclusion

This study demonstrates that AH9 exhibits significant anti-migratory activity in endothelial cells and causes the downregulation of ERK 1/2 phosphorylation. The reduction in ERK phosphorylation by AH9 could be the result of reduced VEGFR expression. Whether this is linked to a reduction in VEGFR phosphorylation requires further investigation. However, as this stage we cannot identify which kinase (MAP3K, MAP2K and MAPK) AH9 is affecting and would need further probing. To add, we cannot rule out that AH9 could be increasing the activity of phosphatases that could cleave off phosphate groups causing the decrease phosphorylation seen. This would also need investigating via phosphatase colorimetric assays for example. As an inhibitor of endothelial cell migration and ERK phosphorylation, AH9 stands as a promising drug candidate for anti-angiogenic therapy and warrants further investigation as a viable option to treat highly vascularised tumours.

## Abbreviations

(HUVECS): Human Umbilical Venous Endothelial Cells
(AH9): 2-bromo-2’5’-dihydroxychalcone
(MTT): 3-(4,5-dimethylthiazol-2-yl)-2,5-diphenyltetrazolium bromide
(VEGF): Vascular Endothelial Growth Factor
(VEGFR-2): Vascular Endothelial Growth Factor Receptor 2
(CVD): Cardiovascular Disease
(AMD): Age Related Macular Degeneration
(ERK 1/2): Extracellular signal-regulated Kinase
(IgG): Immunoglobulin G
(MAPK): Mitogen-activated protein kinase
(FGF): Fibroblast Growth Factor
(FGFR): Fibroblast Growth Factor Receptor
(sRTKI’s): Small Receptor Tyrosine Kinase Inhibitors
(HeLa): Human Cervical Carcinoma Cells
(CAM): Chorioallantoic Membrane Model

