## Supplement Material for "2-bromo-2’5’-dihydroxychalcone analogue Inhibits Endothelial Migration by Targeting VEGF-induced ERK 1/2 Phosphorylation"

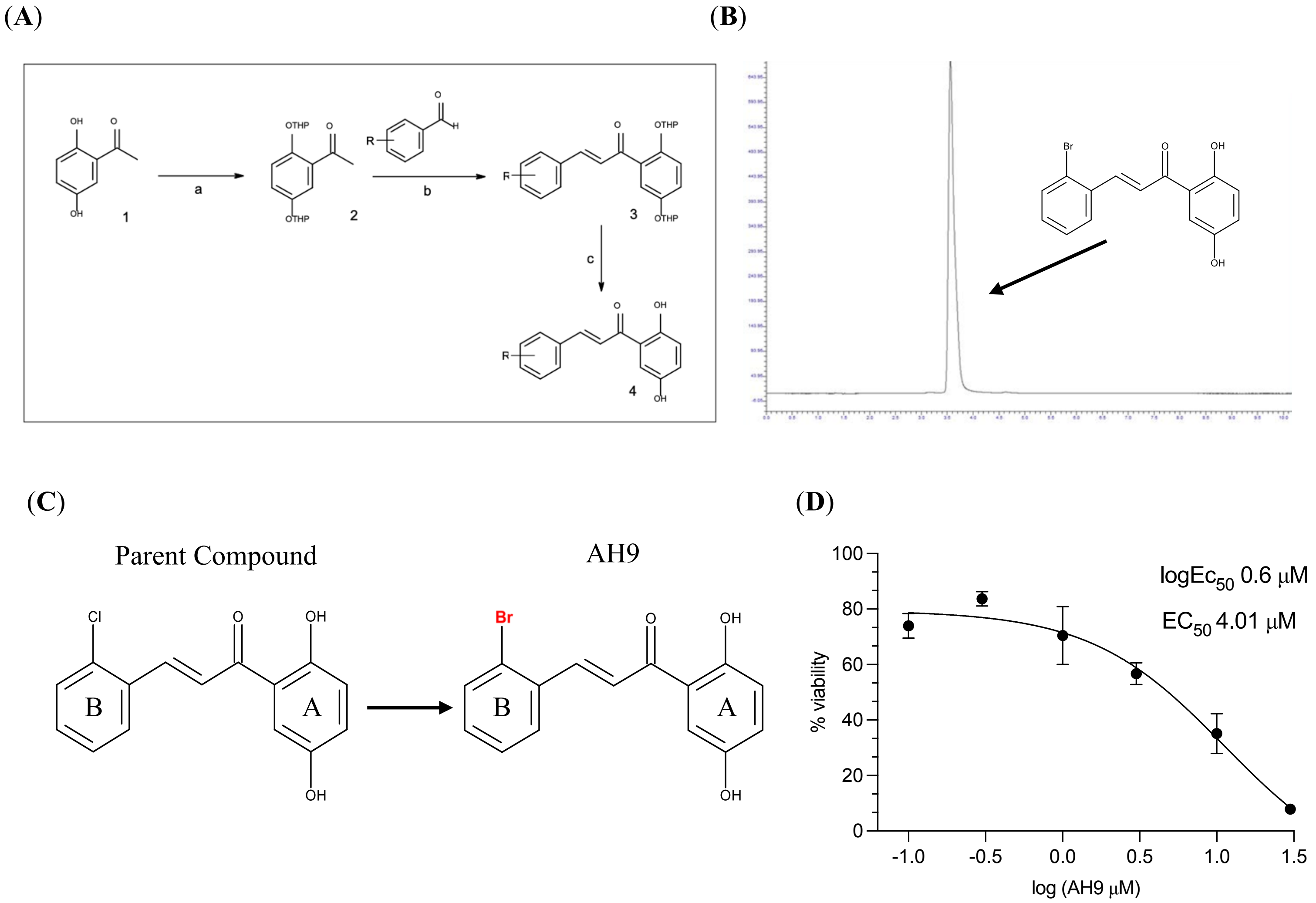


**Figure 1. Chemical structures of parent compound (2-chloro-2’5’-dihydroxchalcone) and AH9 (2-bromo-2’5’-dihydroxychalcone).** Similarities are seen between the two diphenylpropenone structures. The only difference is the halogen on the B ring ortho position from a chlorine to a bromine.

(**A**)

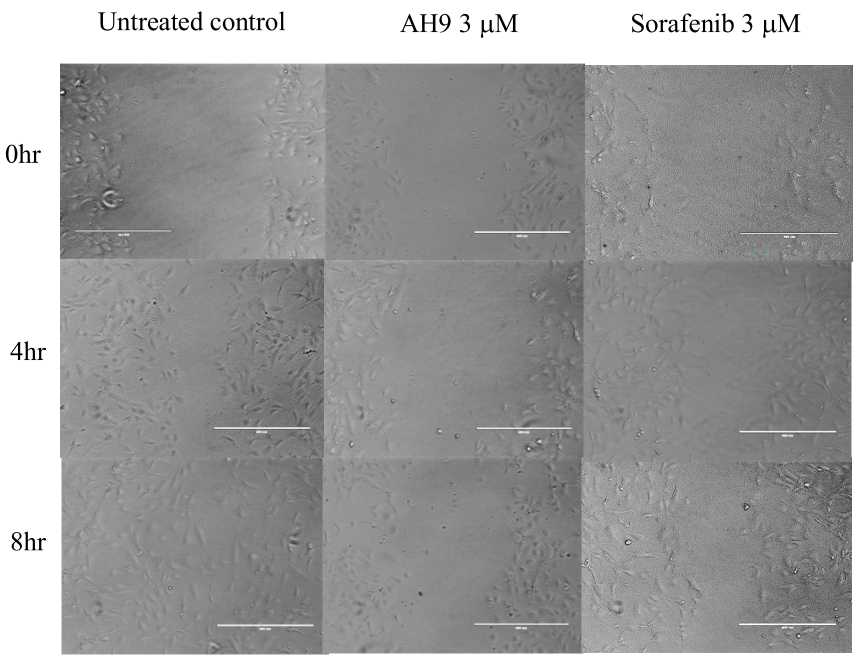


(**B**)

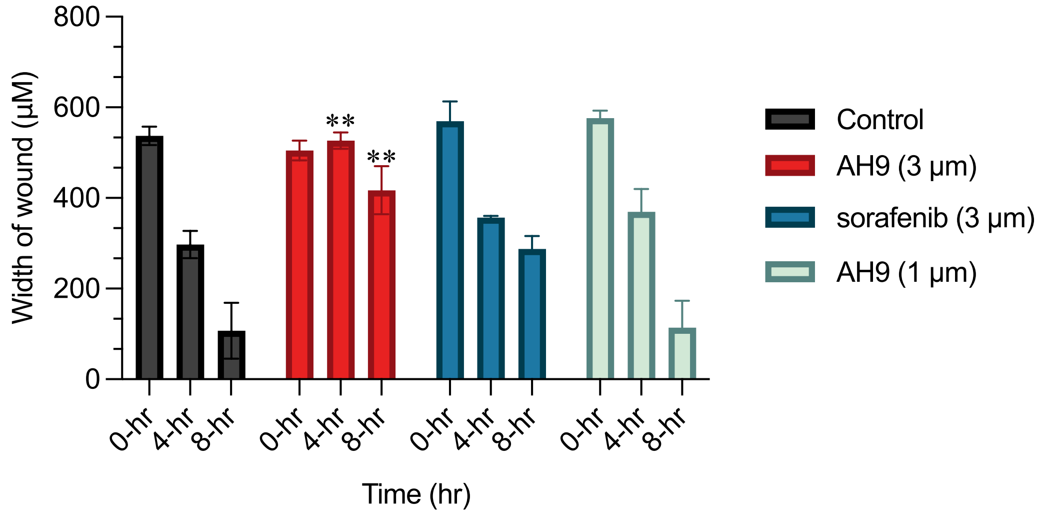


**Figure 2. AH9 inhibited HUVEC migration. A)** HUVECs cultured in 6 well plates were scratched with 1ml pipette tip and treated with different concentration of AH9 and Sorafenib. Images were taken at regular time points (0hr, 4hr and 8hr) using the EVOS FL Cell Imaging System (magnification, X100). **B)** Effects of AH9 and Sorafenib on HUVEC migration at each of the 3 time points (0hr, 4hr and 8hr) were quantified by assessing the width of the wound. Data presented as mean percentage changes vs untreated control (0hr normalised to 100%) of three independent experiments ± SEM. Responses were compared using one way-ANOVA with Tukey-Kramer multiple comparisons post hoc t test (AH9 vs untreated control after 4-hr,

** = (-4.5% p = 0.0023), AH9 vs untreated control at 8-hr, ** = (17.41% p = 0.0079). **C)** Individual width of wound measurements of the effects AH9, Sorafenib and untreated control on HUVEC migration. Data presented as mean micrometres of three independent experiments ± SEM. Responses were compared using one way-ANOVA with Tukey-Kramer multiple comparisons post hoc t test.

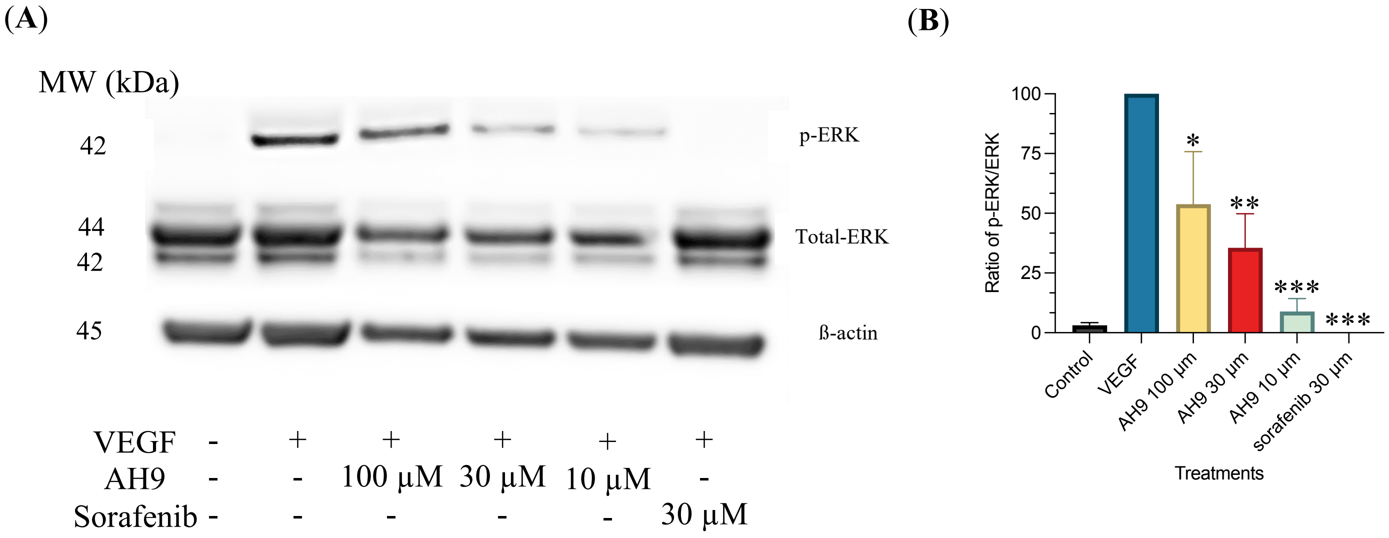


**Figure 3. AH9 inhibited VEGF-induced ERK1/2 phosphorylation. A)** HUVECs were treated with different concentrations of AH9 with or without VEGF (30 ng/ml). Cells were then lysed using laemelli buffer. Protein lysates were then subjected to electrophoresis and transferred onto nitrocellulose membranes. Membrane was then probed for p-ERK 1/2, total ERK ½ and β-actin. **B)** Membrane captures were quantified using densitometry, levels of proteins were quantified and normalised using β-actin as an internal control. Ratio of phosphorylated ERK vs total ERK (normalised to VEGF control) were expressed as means of three independent experiments ± SEM. Responses were compared using one way-ANOVA with Tukey-Kramer multiple comparisons post hoc t test 100 µM (* = (46%, p = 0.003), 30 µM (** = (64%, p = 0.0002) and 10 µM (*** = (91%, p = 0.0001).

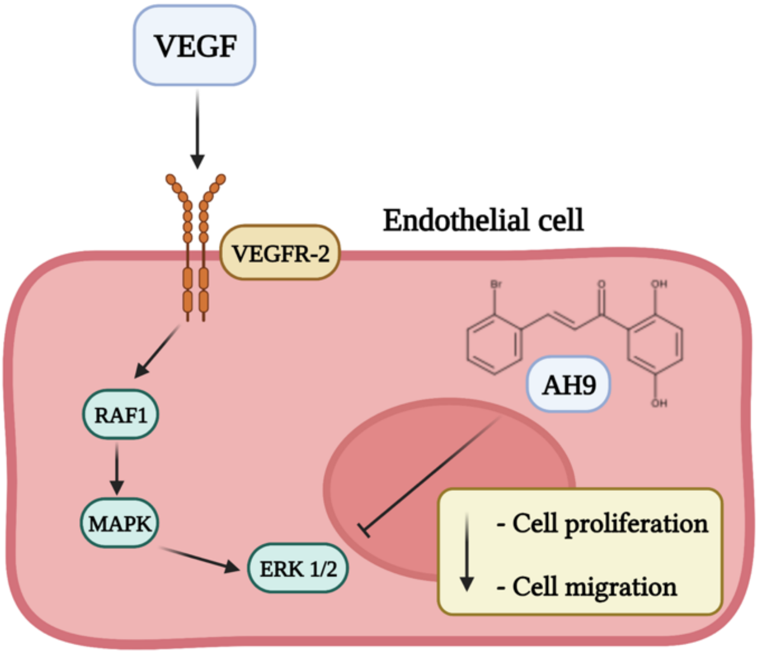


**Figure 4. Potential mechanism of action for AH9 induced ERK1/2 inhibition via the VEGF/VEGFR2 pathway.**
